# LmTag: functional-enrichment and imputation-aware tag SNP selection for population-specific genotyping arrays

**DOI:** 10.1101/2022.01.28.478108

**Authors:** Dat Thanh Nguyen, Quan Hoang Nguyen, Nguyen Thuy Duong, Nam Sy Vo

## Abstract

Despite the rapid development of sequencing technology, single-nucleotide polymorphism (SNP) array is still the most cost-effective genotyping solutions for large-scale genomic research and applications. Recent years have witnessed the rapid development of numerous genotyping platforms of different sizes and designs, but population-specific platforms are still lacking, especially for those in developing countries. We aim to develop methods to design SNP arrays for thse countries, so the arrays should be cost-effective (small size), yet can still generate key information needed to associate genotypes with traits. A key design principle for most current platforms is to improve genome-wide imputation so that more SNPs (imputed tag SNPs) not included in the array can be predicted. However, current tag SNP selection methods mostly focus on imputation accuracy and coverage, but not the functional content of the measured and imputed SNPs. It is those functional SNPs that are most likely associated to traits. Here, we propose LmTag, a novel method for tag SNP selection that not only improves imputation performance but also prioritizes highly functional SNP markers. We apply LmTag on a wide range of populations using both public and in-house whole genome sequencing databases. Our results showed that LmTag improved both functional marker prioritization and genome-wide imputation accuracy compared to existing methods. This novel approach could contribute to the next generation genotyping arrays that provide excellent imputation capability as well as facilitate array-based functional genetic studies. Such arrays are particularly suitable for under-represented populations in developing countries or non-model species, where little genomics data are available while investment in genome sequencing or high-density SNP arrays is limited.

## Introduction

Single-nucleotide polymorphism (SNP) arrays and recent technology whole-genome sequencing (WGS) have been widely used in genomic research and applications. While WGS is attractive due to its ability to capture all genetic variation in the genome, SNP arrays have been the most widely used strategy due to several advantages such as cost-effectiveness, reliability of the technology, and light computational requirement (Tam et al., 2019). SNP arrays still play important role in Genome-wide association studies (GWAS), which have facilitated the detection of DNA variants associated with human complex traits, including disease traits, leading to numerous proven and potential translational applications toward new diagnoses and therapeutics over the last decade (Visscher et al., 2017). However, due to the small number of SNPs that can be included, array-based genomic studies often required imputation to increase the number of variants for association tests by predicting the genotypes at the SNPs that are not directly genotyped in the study samples. The performance of imputation is affected by three main factors, including imputation algorithms (Das et al., 2016), imputation reference panels (Huang et al., 2015; McCarthy et al., 2016), and the design of SNP arrays (Nelson et al., 2013).

Available genomic studies have focused mainly on European descent, accounting for approximately 79% of all GWAS participants, while the overall European population comprises about 16% of the total global population (Martin et al., 2019; Peterson et al., 2019). Given that the majority of human functional genetic variants are population-specific and rare (Nelson et al., 2012; Consortium et al., 2015), the imbalance in current population genetic data resources implies a critical problem. Important variants with low frequencies or completely absent in European populations may be missed by GWAS discoveries so far (Wojcik et al., 2019). Consequently, disease risk predictions, which benefit the clinical arena, are currently restricted in the European ancestry population (Duncan et al., 2019). This is a critical issue, especially for the majority of the world population, who are under-represented in genomic studies. These underrepresented populations include both minority ethnic groups within high-income countries, and citizens of low and middle-income countries (Lewis and Vassos, 2020). This fact leads to an urgent and unmet demand to develop and use customized genotyping platforms for under represented populations (Tam et al., 2019). Indeed, population-specific genotyping arrays such as the UK Biobank Axiom Array (Bycroft et al., 2018), the Axiom-NL Array (Ehli et al., 2017), the TWB Array (Chen et al., 2016), the Axiom China Kadoorie Biobank Array (Dai et al., 2019), the Japonica and Japonica NEO Arrays (Kawai et al., 2015; Sakurai-Yageta et al., 2020), and the Axiom KoreanChip (Moon et al., 2019) have been successfully implemented to facilitate genomic studies in these populations.

To develop such arrays, various strategies to select tag SNPs are employed. A tag SNP is a SNP that can represent a group of SNPs called a haplotype due to strong associations between these neighboring alleles (known as linkage disequilibrium, LD). Tag SNP selection methods can be classified into two main categories including block-based (Johnson et al., 2001; Patil et al., 2001; Sebastiani et al., 2003), and LD-based approaches (Carlson et al., 2004; Liu et al., 2010; Hoffmann et al., 2011a; Wojcik et al., 2018). The former approach involves partitioning the whole chromosome into blocks, often relying on a predefined haplotype block structures or simply based on genomic distance. For example, in the early generation of human genotyping SNP array, tag SNPs were selected at intervals of approximately each 5-kilobase with a minor allele frequency of at least 5% (Gibbs et al., 2003). This strategy has also been widely adopted in animal genetics, commonly referred to as the equidistance method (Shashkova et al., 2020; Herry et al., 2018). On the other hand, the latter approach utilizes LDs among nearby SNPs to find tag SNPs with a greedy approach to maximize LD coverage (Weale et al., 2003; Sakurai-Yageta et al., 2020; Wojcik et al., 2018). A typical algorithm starts with a set of targeted SNPs, then weights each SNP candidate by the number of neighbor SNPs (within a specific genomic distance) that have pairwise LD *r*^2^ greater than or equal to a specific threshold, e.g., 0.8. The SNP with the highest score is then selected, and the associated SNPs are removed from the targeted set. These steps are iterated until reaching the desired number of tag SNPs or no more SNP satisfies the LD *r*^2^ threshold (Carlson et al., 2004; Weale et al., 2003). In addition, multi-marker LD approach (Wang and Jiang, 2008; Hao, 2007; Liu et al., 2010), pairwise LD hybrid tag SNP selection (Hoffmann et al., 2011a),cross-population prioritizing scheme (Wojcik et al., 2018) also aimed to improve LD coverage and imputation accuracy. Despite the efforts, these strategies still have certain limitations. Firstly, it is unclear that tag SNP selection approaches to maximize LD coverage or genomic distance can provide the best imputation accuracy performance, which is the golden standard of SNP array assessment nowadays (Nelson et al., 2013; Wojcik et al., 2018). Secondly, SNPs on genotyping arrays are typically not causal variants because they are chosen to be highly LD correlated with neighboring SNPs to cover large genomic regions to allow for imputing unmeasured SNPs, a common design practice in the greedy paradigm (Schaid et al., 2018).

To address these challenges, we introduce a novel method called LmTag, which faciliates design of functional-enrichment, imputation-aware, and population-specific SNP arrays. Firstly, LmTag uses a robust statistical modeling to systematically integrate LD information, minor allele frequency (MAF), and physical distance of SNPs into the imputation accuracy score to improve tagging efficiency. Secondly, LmTag adapts the beam search framework (Lowerre, 1976) to prioritize both variant imputation scores and functional scores to solve the tag SNP selection problem. We apply LmTag and comprehensively compare it with common approaches of tag SNP selection using a wide range of both public and in-house genomics datasets. Our benchmarking results suggest that LmTag improves both imputation performance and prioritization of functional variants. Furthermore, we show that tagging efficiency of tag SNP sets selected by LmTag are sustainability higher than existing genotyping arrays, indicating the potential improvements for future genotyping platforms.

## Results

### Overview of LmTag pipeline

An overview of LmTag is presented in Figure 1. The method includes three key steps: (i) Imputation accuracy modeling, (ii) Functional scoring, and (iii) Functional tag SNP selection. In the first step, a theoretical array (set of tag SNPs) is simulated, and imputation accuracy scores of the corresponding tagged SNPs are estimated by leave-one-out cross-validation (details in the method section). A linear model is then employed to assess imputation accuracy scores of tagged SNPs based on pairwise LD *r*^2^, MAF of tag SNPs (those included in the array), MAF of tagged SNPs (not included in the array), and distances between tag SNPs and tagged SNPs. In the second step, SNPs are functionally scored based on public databases including the GWAS catalog (MacArthur et al., 2017), the ClinVar (Landrum et al., 2018), and the Combined Annotation-Dependent Depletion (CADD) (Kircher et al., 2014) to enrich functional variants in the array design. Finally, parameters from the model are used to estimate imputation accuracy score for each SNP. These estimated scores, together with the functional ranking of SNPs, are then used in functional-enrichment tag SNP selection by the beam search algorithm with beam width parameter *K* (Lowerre, 1976). Further details are described in the “Methods” section.

**Figure 1:**
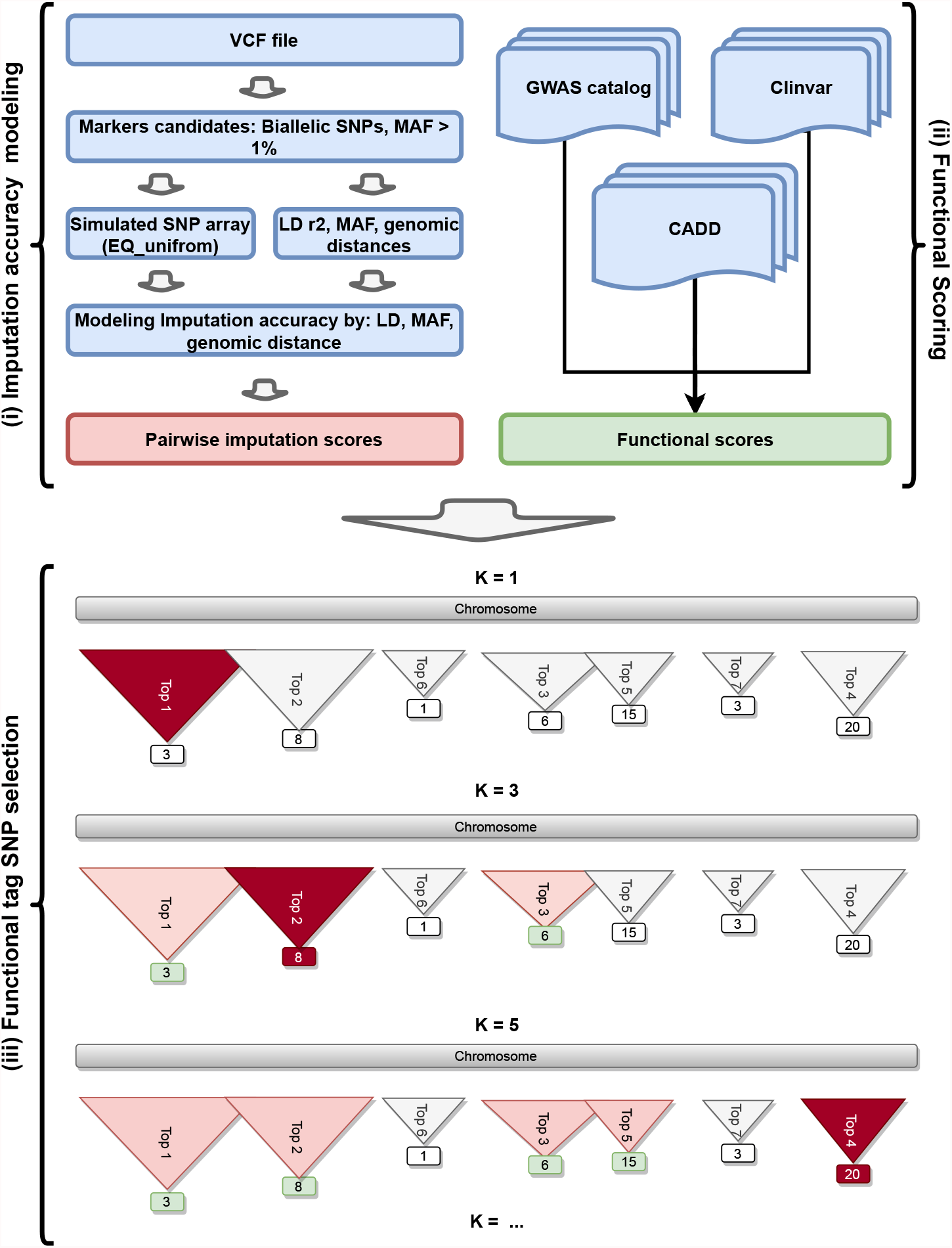
Overview of LmTag. (i) Imputation accuracy modeling, this includes modeling imputation accuracy metric as a function of LD, MAFs, and genomic distances. (ii) Functional scoring, this includes steps of weighting functional scores of SNPs based on public databases. (iii). Functional tag SNP selection, Imputation capability of each SNP is represents as triangles while functional scores are showed in the lower rectangles. When *K* = 1, the beam search algorithm becomes the best-fist search that select SNP with highest estimated imputation performance - colored bold red triangles. When *K >* 1, the algorithm select top *K* SNPs with the highest estimated imputation performances - colored light pink triangles, the functional scores in these SNPs - colored light green is weighted to find the highest functional SNPs as tag SNPs - colored bold red triangles.

### LmTag improves functional enrichment in tag SNP selection

LmTag performancce is compared against commonly used tag SNP selection methods including TagIt (Weale et al., 2003), FastTagger (Liu et al., 2010), EQ_uniform (Shashkova et al., 2020), and EQ_MAF (Herry et al., 2018) by two main metrics: functional enrichment and imputation performance. The benchmarking is performed in both in-house and public genomics datasets including pilot phase data from the 1000 Vietnamese Genomes Project (1KVG), and data of three super populations comrising obtained from the 1000 Genomes Project samples re-sequenced by New York Genome Center (1KGP-NYGC) (Byrska-Bishop et al., 2021). Overall, four populations including Vietnamese pilot phase (VNP), East Asian (EAS), European (EUR), and South Asian (SAS) comprising WGS data of 504, 504, 503, and 489 individuals respectively are included in the analysis. Further details of datasets and metrics used in compassion experiments are described in the “Methods” section.

The summary results of functional enrichment in tag SNP selection of LmTag, EQ_uniform, EQ_MAF, TagIt, FastTagger, and baseline (mean functional score and proportion of biological evidenced markers in all SNPs in the population) are reported in Tables 3, 4 and visualized in Figures 2, 3. LmTag is evaluated with various beam width parameters *K*=1, 10, 20, 30, 50, 100, 200, denoted as LmTag_K1, LmTag_K10, LmTag_K30, LmTag_K50, LmTag_K100, LmTag_K200, respectively. The results are collected from all four populations EAS, EUR, SAS, and VNP under the 32,000 tag SNPs setting.

**Table 1:** Datasets used in this study.

| Populations | Number of samples | Total markers | GWAS markers | Clinvar markers |
| --- | --- | --- | --- | --- |
| VNP | 504 | 382700 | 5064 | 1590 |
| EAS | 504 | 405234 | 5160 | 1617 |
| SAS | 489 | 486024 | 5868 | 1876 |
| EUR | 503 | 456166 | 6168 | 1814 |

**Table 2:**
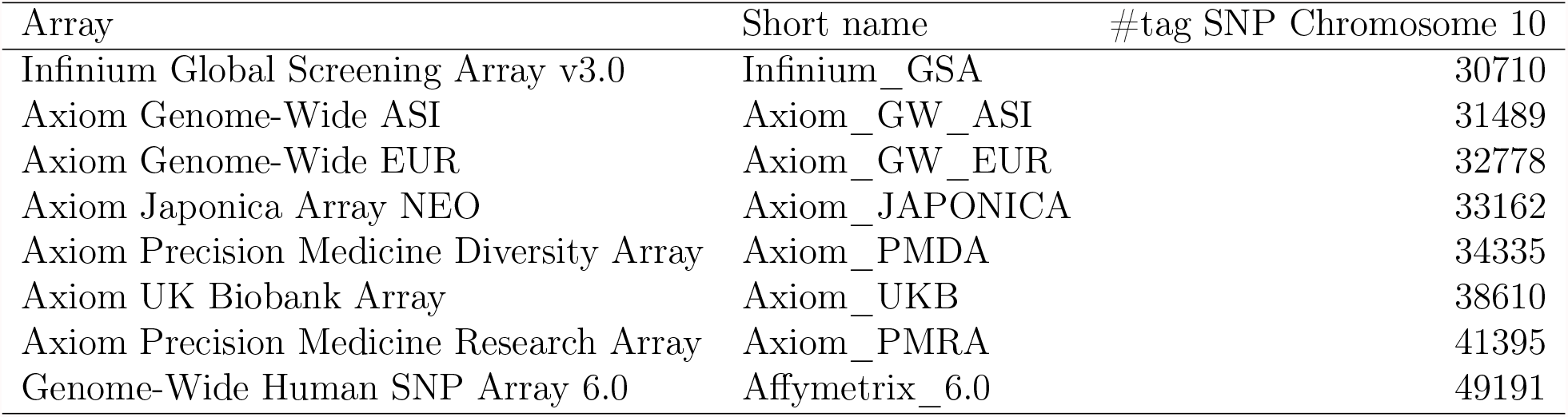
Genotyping arrays used in this study.

**Table 3:** Mean CADD scores in ‘PHRED-scaled’ and its equivalent percentile of tagSNP selected by LmTag (with *K* = 1, 10, 20, 30, 50, 100, 200), EQ_uniform, EQ_MAF, TagIt and FastTagger. Control shows mean CADD scores in ‘PHRED-scaled’ and its equivalent percentiles of all input markers by each population.

| Methods | EAS |  | EUR |  | SAS |  | VNP |  |
| --- | --- | --- | --- | --- | --- | --- | --- | --- |
|  | Mean | Mean | Mean | Mean | Mean | Mean | Mean | Mean |
|  | CADD | percentile | CADD | percentile | CADD | percentile | CADD | percentile |
| Baseline | 2.95 | 38.65 | 2.96 | 38.77 | 2.96 | 38.74 | 2.91 | 38.20 |
| EQ_uniform | 2.96 | 38.64 | 2.96 | 38.70 | 2.95 | 38.74 | 2.89 | 38.03 |
| EQ_MAF | 3.03 | 38.73 | 3.00 | 38.58 | 2.99 | 38.38 | 2.96 | 38.11 |
| TagIt | 2.96 | 38.23 | 2.92 | 38.02 | 2.99 | 38.52 | 2.93 | 37.99 |
| FastTagger | 3.04 | 38.81 | 2.94 | 38.63 | 2.96 | 38.83 | 3.04 | 39.34 |
| LmTag_K1 | 2.96 | 38.33 | 2.93 | 38.19 | 2.92 | 38.29 | 2.92 | 37.97 |
| LmTag_K10 | 3.63 | 45.51 | 3.58 | 45.22 | 3.46 | 44.24 | 3.57 | 44.83 |
| LmTag_K20 | 3.84 | 47.55 | 3.76 | 47.02 | 3.69 | 46.49 | 3.79 | 46.97 |
| LmTag_K30 | 3.98 | 48.77 | 3.93 | 48.68 | 3.86 | 48.08 | 3.93 | 48.25 |
| LmTag_K50 | 4.20 | 50.76 | 4.16 | 50.66 | 4.11 | 50.33 | 4.12 | 49.91 |
| LmTag_K100 | 4.54 | 53.39 | 4.51 | 53.59 | 4.48 | 53.45 | 4.45 | 52.67 |
| LmTag_K200 | 4.82 | 55.65 | 4.87 | 56.48 | 4.86 | 56.44 | 4.77 | 55.20 |

**Table 4:** Percentages of GWAS and Clinvar makers covered by 32000 tag SNPs selected by LmTag (with *K* = 1, 10, 20, 30, 50, 100, 200), EQ_uniform, EQ_MAF, TagIt and FastTagger over total number of GWAS and Clinvar makers in each population. Baseline shows the relative proportion of GWAS and ClinVar markers under the uniform distributed in tag SNP selection, i.e,. baseline values are computed as 32000 devide for total number of marker in the examined populations.

| Methods | EAS |  | EUR |  | SAS |  | VNP |  |
| --- | --- | --- | --- | --- | --- | --- | --- | --- |
|  | GWAS | Clinvar | GWAS | Clinvar | GWAS | Clinvar | GWAS | Clinvar |
| Baseline | 7.90 | 7.90 | 7.01 | 7.01 | 6.58 | 6.58 | 8.36 | 8.36 |
| EQ_uniform | 7.67 | 8.16 | 6.94 | 7.94 | 6.56 | 6.82 | 8.20 | 8.18 |
| EQ_MAF | 15.64 | 10.58 | 15.26 | 8.82 | 15.80 | 8.74 | 16.17 | 11.45 |
| TagIt | 10.00 | 9.96 | 9.48 | 9.54 | 9.34 | 9.28 | 9.91 | 10.50 |
| FastTagger | 7.83 | 7.24 | 6.84 | 8.88 | 6.56 | 7.36 | 8.06 | 10.06 |
| LmTag_K1 | 9.34 | 8.60 | 8.20 | 8.88 | 8.11 | 7.68 | 9.06 | 8.81 |
| LmTag_K10 | 11.61 | 10.51 | 10.42 | 10.36 | 9.90 | 8.96 | 11.02 | 11.07 |
| LmTag_K20 | 12.83 | 11.94 | 10.85 | 10.97 | 10.77 | 9.59 | 12.03 | 11.70 |
| LmTag_K30 | 13.76 | 12.68 | 11.54 | 11.36 | 11.32 | 10.23 | 12.58 | 12.77 |
| LmTag_K50 | 14.79 | 13.30 | 12.42 | 12.02 | 12.05 | 11.25 | 13.88 | 14.15 |
| LmTag_K100 | 17.27 | 15.71 | 13.85 | 14.28 | 13.97 | 13.01 | 15.86 | 16.16 |
| LmTag_K200 | 19.96 | 16.51 | 16.05 | 16.32 | 15.61 | 14.71 | 18.13 | 18.05 |

**Figure 2:**
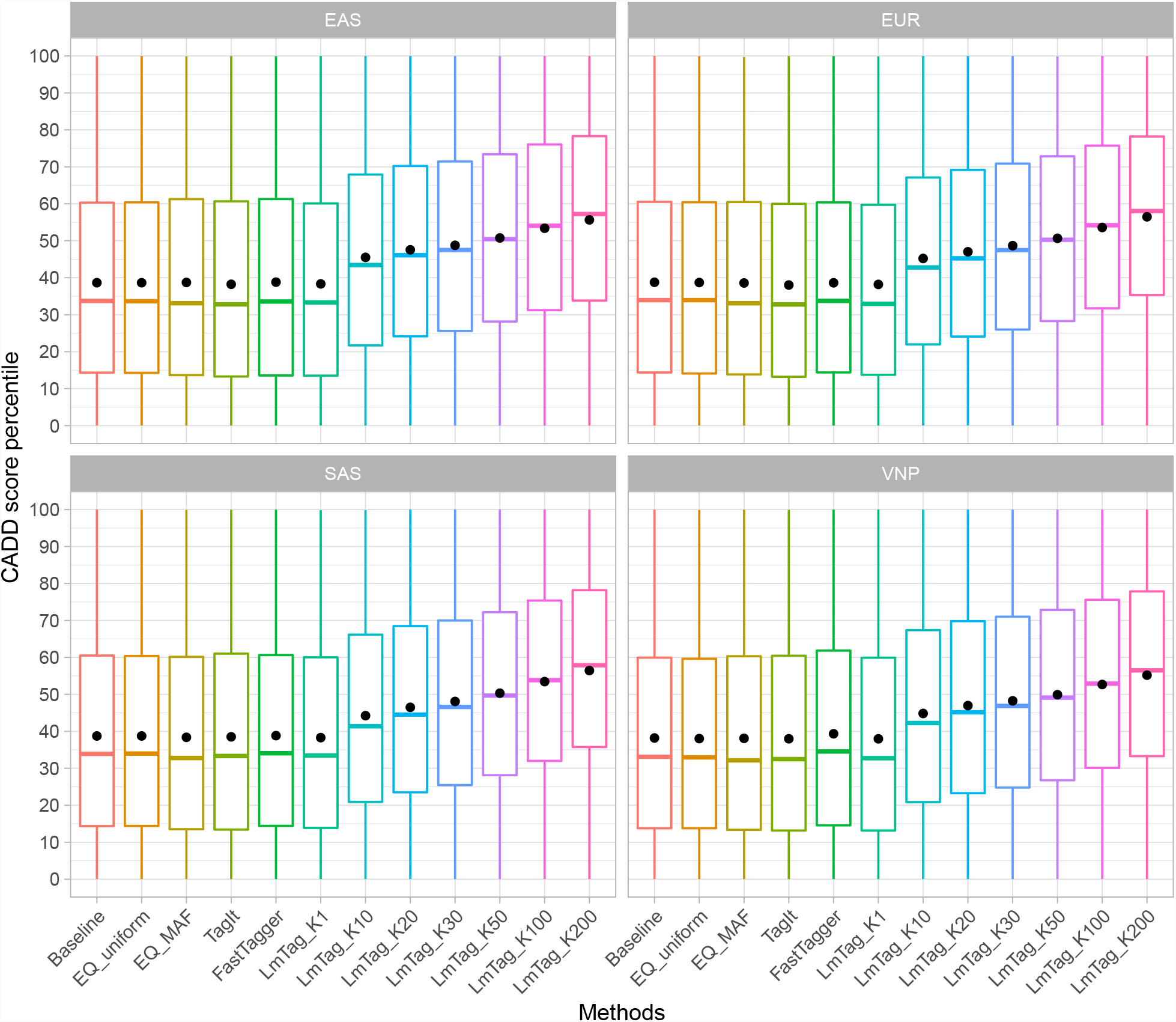
Mean percentile of CADD scores of tag SNP selected by LmTag (with K=1, 10, 20, 30, 50, 100, 200), EQ_uniform, EQ_MAF, TagIt and FastTagger. Baseline shows mean percentile of CADD scores of all input markers (32,000 SNPs) in each population.

**Figure 3:**
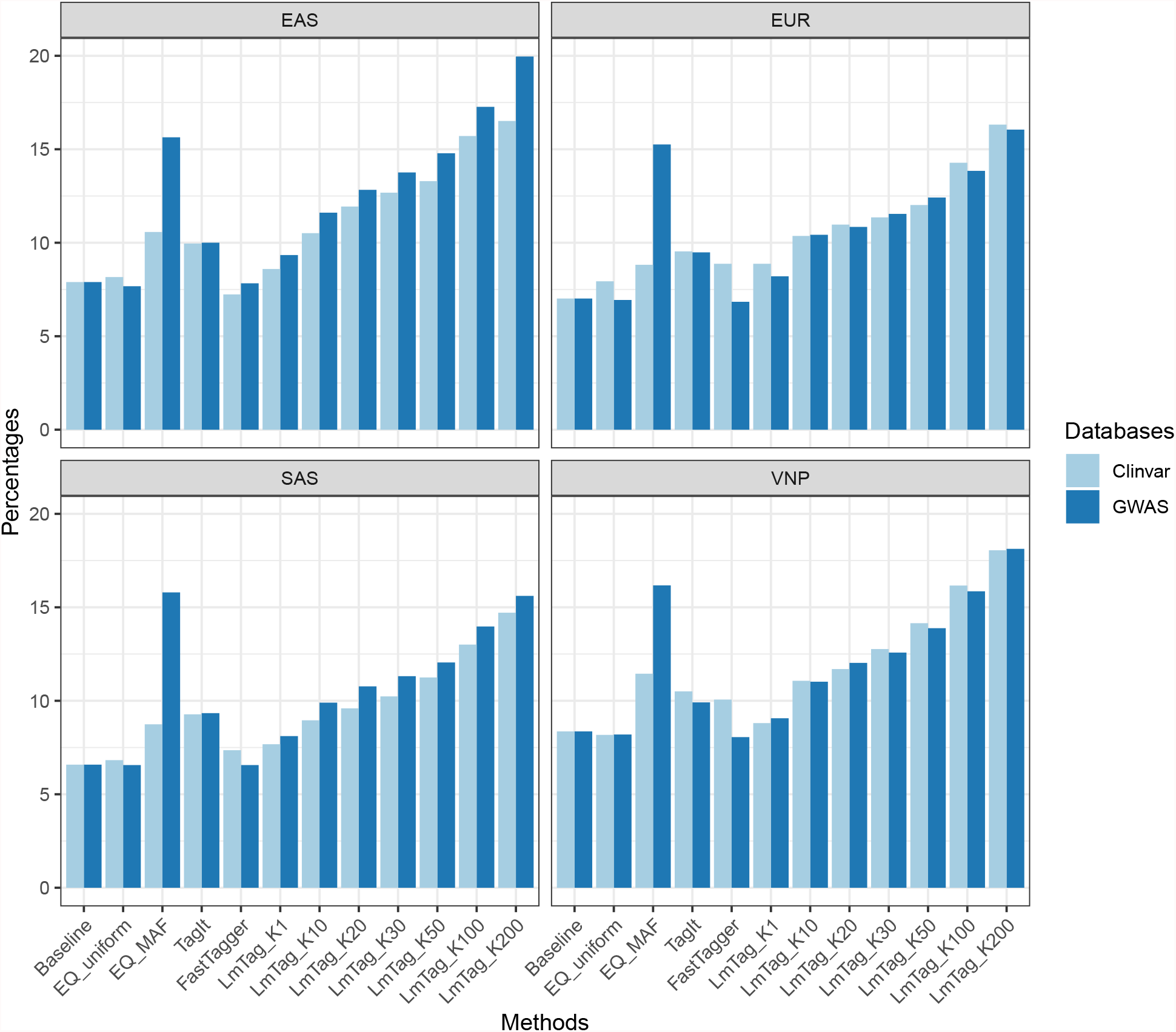
Percentages of GWAS and Clinvar makers covered by 32000 tag SNPs selected by LmTag (with *K* = 1, 10, 20, 30, 50, 100, 200), EQ_uniform, EQ_MAF, TagIt and Fast-Tagger over total number of GWAS and Clinvar makers in each population. Baseline shows percentages of GWAS and ClinVar markers covered over total number of GWAS and Clinvar makers in each population corresponding 32000 tag SNP scaffold.

In general, LmTag shows a significant improvement in functional prioritization with almost zero imputation performance trade-off. Particularly, in comparison to the baseline and other methods, LmTag (at *K* = 200) obtains significant improvements with approximately 2-fold enrichment in terms of selection GWAS and ClinVar markers; and yet increases averagely 15-17% CADD score percentile ranking in term of selection population-wide variants as tag SNPs.

When *K* is set as 1, LmTag becomes a standard greedy algorithm with the “best-first” search approach, i.e., no optimization is applied for selecting functional variants. In this setting, mean CADD scores, mean CADD percentiles, proportions of GWAS, and ClinVar markers selected by LmTag are comparable with the baseline and other methods, as expected. The mean CADD scores of tag SNPs selected by LmTag_K1 vary from 2.92 to 2.96 across examined populations, and are in the same range with the baseline, which varies from 2.91 to 2.96. Other methods also yield comparable performances with LmTag_K1, ranging from 2.89 to 3.04. Conversion from ‘PHRED-scaled’ score into percentile scale shows mean CADD score percentile of LmTag_K1 and the others are equivalent with the rank from 37.97 to 39.34. In other words, under the setting of no optimization for functional SNPs, CADD scores / percentiles of tag SNP distribute equivalently regardless of the method of choice. Similarly, when considering prioritization of markers using biological evidence databases, the proportions of GWAS and ClinVar marker selected by LmTag_K1, and other methods are mostly comparable to the baseline except for GWAS marker proportions of EQ_MAF as shown in Figure 3. Under the baseline scenario, the expected proportions of GWAS and ClinVar in 32,000 tag SNPs are 7.90%, 7.01%, 6.58%, and 8.36% in EAS, EUR, SAS, and VNP, respectively. The corresponding ranges for LmTag_K1, EQ_uniform, TagIt and FastTagger are 7.68 - 9.34%, 6.58 - 8.36%, 9.34 - 10.50%, respectively. Notably, the EQ_MAF method appeared to select slightly higher proportions of ClinVar markers, from 8.74-11.45%, and significantly more GWAS markers ranging from 15.26 to 16.17% that are possibly explained by the detection power is bias toward high frequency variant in both clinical and association studies.

When the value of *K* increase, as expected, a clear improvement of functional enrichment is shown as detailed in Tables 3, 4, 5 and Figures 2, 3. Consistently, CADD scores and proportions of GWAS and ClinVar show a strong positive correlation with the increase of *K*, while the overall imputation accuracy is converged or experienced very small changes. For example, in the VNP population, the overall imputation accuracies stay stable around 89.80% despite the dramatic changes of *K* values from 1 to 200. While the functional SNP prioritization process do not reduce LmTag imputation performance, there are significant improvements in tag SNP functional scores. The mean CADD score percentile increase from 37.97 to 44.83, 46.97, 48.25, 49.91, 52.67, and 50.20 in response to *K* value increasing from 1 to 10, 20, 30, 50, 100, and 200, respectively. It is noted that, mean CADD score percentile values are computed by taking the average percentile ranks of all selected tag SNPs and not by directly converting from the mean of CADD “PHRED-scaled” scores. Importantly, the GWAS and ClinVar proportions covered by 32000 tag SNP also increase more than 2-folds, both from 8.36% to 18.13%, and 18.05%, respectively. Consistent improvements could be observed clearly in other populations. At K=200, the average CADD score mean percentiles of EAS, EUR, SAS increase to 55.65, 56.48, 56.44, respectively. The number of GWAS and ClinVar markers selected are also significantly improved in these populations.

**Table 5:**
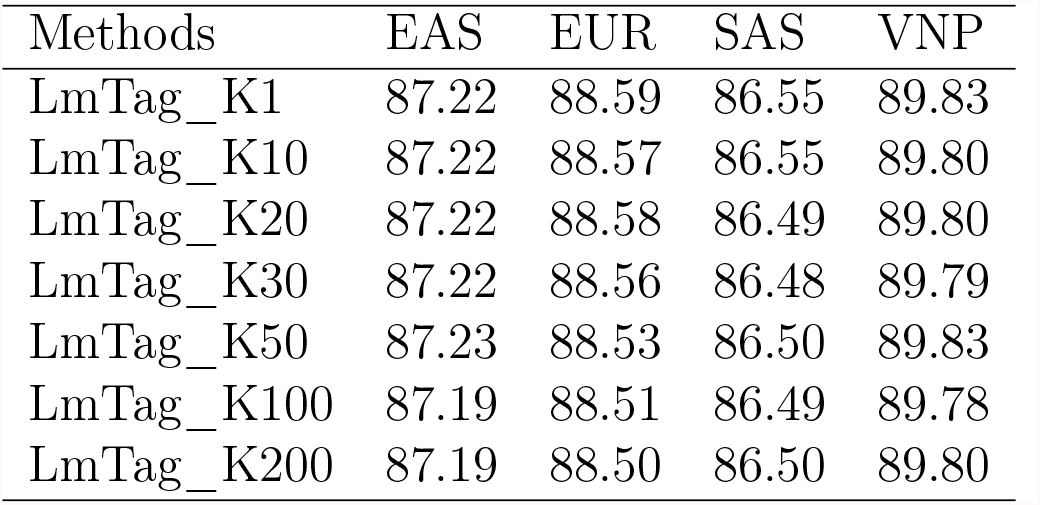
Overall imputation accuracies (mean imputation *r*^2^ of all markers) of LmTag at 32000 tag SNP scaffold with various K values (*K* = 1, 10, 20, 30, 50, 100, 200).

### LmTag demonstrates superior tagging efficiency

Regarding imputation performance, LmTag outperforms other methods in both imputation accuracy and imputation coverage. The *K* parameter used in this comparison is 200, while the number of tag SNPs is set at various cutoffs. Details are reported in Table 6, and Figure 4. Regarding imputation accuracy, LmTag is the top performer, followed by TagIt, and EQ_uniform while the worst performers are interchanged between EQ_MAF and FastTagger depending on population. At the cutoff of 32,000, performance differences are substantially large between LmTag against EQ_uniform, EQ_MAF, and FastTagger but smaller against TagIt. For example, in the EAS population, LmTag obtains 87.19% overall imputation accuracy compared with 86.29%, 82.51%, 82.33%, and, 78.10% achieve by TagIt, EQ_uniform, FastTagger, and EQ_MAF respectively. The same trend is also observed in EUR, SAS, and VNP with 88.50%, 86.50%, and 89.80% imputation accuracies achieved by LmTag_200. In terms of imputation coverage, LmTag also produces the highest performance. Taking imputation *r*^2^ threshold of 80% as an example, LmTag yields the imputation coverage of 83.65%, 85.25%, 81.66%, and 87.81% in EAS, EUR, SAS and VNP while the second-ranked performer obtains 82.11%, 84.08%, 80.13%, and 87.04% respectively.

**Table 6:** Overall imputation accuracies (mean imputation *r*^2^ of all markers), and imputation coverages (proportions of markers with imputation *r*^2^ greater than or equal to 0.8 over total marker in population) for each population corresponding to multiple cutoffs ranging from 8000 to 32000 tag SNP selected by LmTag (with *K* = 200), EQ_uniform, EQ_MAF, TagIt and FastTagger.

| # tag SNP | Methods | EAS |  | EUR |  | SAS |  | VNP |  |
| --- | --- | --- | --- | --- | --- | --- | --- | --- | --- |
| | | Mean imputation $r^2$ | Imputation coverage | Mean imputation $r^2$ | Imputation coverage | Mean imputation $r^2$ | Imputation coverage | Mean imputation $r^2$ | Imputation coverage |
| 32000 | LmTag_K200 | 87.19 | 83.65 | 88.50 | 85.25 | 86.50 | 81.66 | 89.80 | 87.81 |
|  | EQ_uniform | 82.51 | 74.17 | 84.67 | 77.01 | 82.10 | 72.15 | 86.03 | 79.61 |
|  | EQ_MAF | 78.10 | 62.91 | 82.19 | 71.00 | 78.73 | 63.32 | 82.78 | 70.98 |
|  | TagIt | 86.29 | 82.11 | 87.71 | 84.08 | 85.71 | 80.13 | 89.23 | 87.04 |
|  | FastTagger | 82.33 | 71.07 | 81.86 | 72.06 | 79.67 | 68.40 | 80.90 | 69.96 |
| 28000 | LmTag_K200 | 86.03 | 81.72 | 87.60 | 83.68 | 85.30 | 79.51 | 88.91 | 86.04 |
|  | EQ_uniform | 81.31 | 72.08 | 83.59 | 74.81 | 80.70 | 69.50 | 84.93 | 77.36 |
|  | EQ_MAF | 76.35 | 58.96 | 80.69 | 67.46 | 76.94 | 59.20 | 81.29 | 67.29 |
|  | TagIt | 84.99 | 80.00 | 86.86 | 82.53 | 84.64 | 78.13 | 88.13 | 85.10 |
|  | FastTagger | 81.41 | 69.31 | 80.68 | 69.83 | 78.15 | 65.28 | 79.19 | 66.86 |
| 24000 | LmTag_K200 | 84.73 | 79.54 | 86.27 | 81.29 | 83.96 | 77.02 | 87.84 | 83.97 |
|  | EQ_uniform | 79.64 | 68.50 | 82.09 | 71.91 | 78.97 | 65.84 | 83.55 | 74.55 |
|  | EQ_MAF | 73.84 | 53.33 | 78.99 | 63.37 | 74.94 | 54.68 | 79.30 | 62.31 |
|  | TagIt | 83.68 | 77.59 | 85.51 | 80.14 | 83.24 | 75.56 | 86.95 | 82.68 |
|  | FastTagger | 80.38 | 67.51 | 78.00 | 65.15 | 75.60 | 61.23 | 77.62 | 64.04 |
| 20000 | LmTag_K200 | 82.99 | 76.54 | 84.67 | 78.43 | 82.12 | 73.91 | 86.41 | 81.31 |
|  | EQ_uniform | 77.54 | 64.77 | 80.09 | 67.55 | 76.81 | 61.56 | 81.85 | 71.00 |
|  | EQ_MAF | 70.80 | 46.89 | 76.34 | 56.49 | 72.02 | 47.86 | 76.82 | 56.10 |
|  | TagIt | 82.02 | 74.65 | 83.95 | 77.01 | 81.34 | 72.04 | 85.60 | 80.06 |
|  | FastTagger | 78.77 | 64.17 | 76.13 | 62.07 | 73.57 | 58.14 | 76.11 | 61.31 |
| 16000 | LmTag_K200 | 80.63 | 72.65 | 82.48 | 74.25 | 79.55 | 69.20 | 84.44 | 77.69 |
|  | EQ_uniform | 74.59 | 58.88 | 77.39 | 62.11 | 73.50 | 55.46 | 79.18 | 65.54 |
|  | EQ_MAF | 66.73 | 38.97 | 73.13 | 48.33 | 68.24 | 39.79 | 73.40 | 47.84 |
|  | TagIt | 79.62 | 70.26 | 81.73 | 72.75 | 78.78 | 67.25 | 83.55 | 76.29 |
|  | FastTagger | 75.18 | 55.70 | 72.28 | 56.27 | 69.62 | 52.34 | 71.87 | 54.47 |
| 12000 | LmTag_K200 | 77.17 | 66.35 | 78.94 | 67.73 | 75.36 | 61.74 | 81.30 | 71.81 |
|  | EQ_uniform | 70.64 | 51.85 | 73.06 | 53.65 | 68.75 | 46.39 | 75.06 | 57.16 |
|  | EQ_MAF | 61.04 | 28.83 | 68.67 | 38.49 | 62.85 | 29.49 | 68.45 | 36.88 |
|  | TagIt | 76.14 | 64.21 | 78.26 | 66.19 | 74.53 | 59.84 | 80.50 | 70.42 |
|  | FastTagger | 72.28 | 50.25 | 67.46 | 49.93 | 64.44 | 45.62 | 67.98 | 48.72 |
| 8000 | LmTag_K200 | 70.67 | 55.37 | 72.45 | 56.67 | 67.79 | 50.25 | 75.18 | 60.83 |
|  | EQ_uniform | 62.80 | 39.60 | 65.79 | 41.52 | 60.25 | 34.14 | 67.96 | 44.41 |
|  | EQ_MAF | 53.16 | 18.84 | 61.29 | 25.03 | 54.76 | 18.12 | 61.11 | 24.52 |
|  | TagIt | 69.48 | 52.56 | 71.88 | 55.23 | 66.96 | 48.07 | 74.40 | 59.01 |
|  | FastTagger | 65.57 | 37.27 | 60.24 | 41.38 | 56.54 | 37.21 | 59.55 | 39.37 |

**Figure 4:**
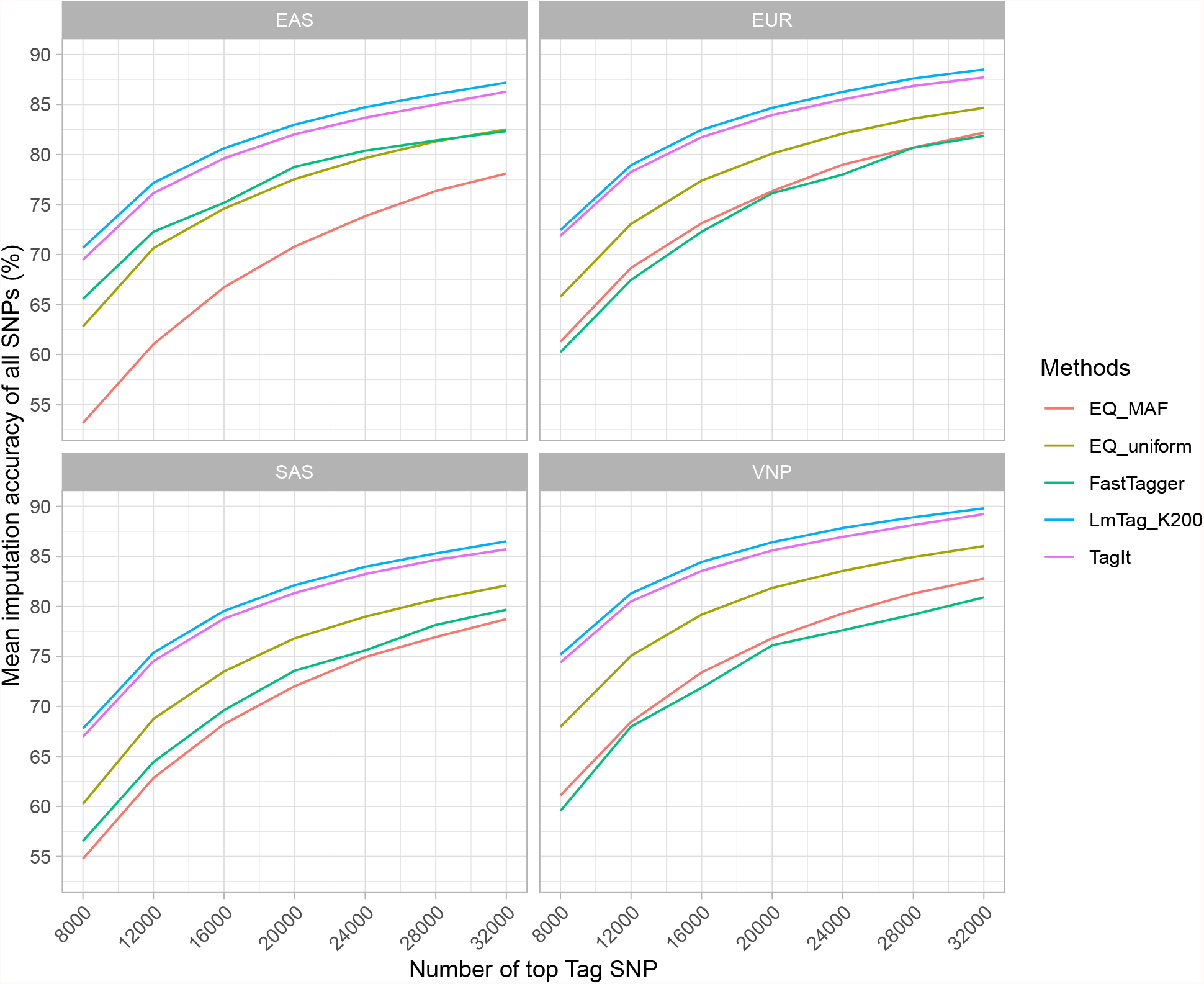
Overall imputation accuracies (mean imputation *r*^2^ of all markers) for each population corresponding to multiple cutoffs ranging from 8000 to 32000 tag SNPs selected by LmTag (with *K* = 200), EQ_uniform, EQ_MAF, TagIt and FastTagger.

To examine potential effects of the number of selected tag SNPs on imputation accuracy and imputation coverage, we further evaluate overall imputation accuracy across different scaffolds by selecting top-ranked SNPs from each population with various cutoffs: 32,000, 28,000, 24,000, 20,000, 16,000, 12,000, and 8,000. Details of overall imputation accuracies are reported in Table 6. We observe that the imputation accuracy and imputation coverage increase in response to the increased number of tag SNPs selected. However, the relationship is not linear as shown in Figure 4 and Figure 5. Nevertheless, LmTag consistently outperforms other methods across all settings. In general, the increasing rates of imputation accuracy and imputation coverage are lower when the numbers of tag SNP is high. In other words, when the scaffolds of the SNP array contain a large enough number of SNPs, adding more tag SNPs do not significantly improve imputation accuracy and imputation coverage compared to those with small scaffolds. For example, adding 4,000 tag SNPs at 12,000 tag SNPs scaffold yield approximately 8% improvement in imputation accuracy compared to the scaffolds of 8,000 SNPs regardless of the method of choice. Meanwhile, increasing 4,000 SNPs to the scaffold of 28,000 results in less than 2% improvement in imputation accuracy. Interestingly, we observe that imputation coverages of all methods dramatically change in response to number of tag SNPs. For example, LmTagK_200 obtains more than 80% coverage with imputation cutoff at 80% at 32,000 tag SNP. The coverage reduces significantly to 50-60% when number of tag SNPs is 8000, and even lower for EQ_MAF to 18-25%.

**Figure 5:**
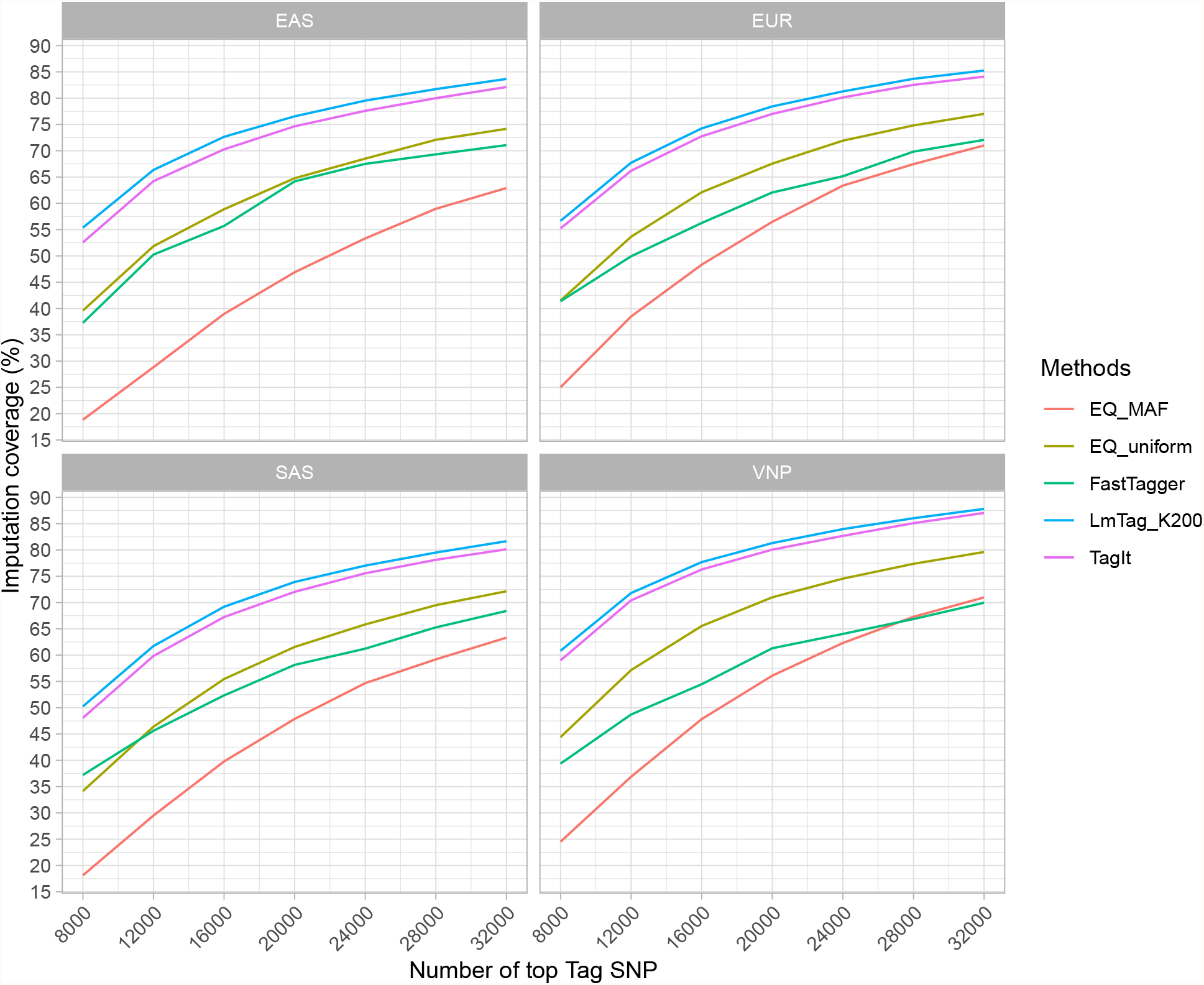
Imputation coverages (proportions of markers with imputation *r*^2^ greater than or equal to 0.8 over total markers in the population) for each population corresponding to multiple cutoffs ranging from 8000 to 32000 tag SNP selected by LmTag (with *K* = 200), EQ_uniform, EQ_MAF, TagIt and FastTagger.

### LmTag helps improve current genotyping arrays

To further explore potential applications of LmTag in designing genotyping arrays. We also compare imputation performances of tag SNPs selected by LmTag (28,000, and 32,000 tag SNPs scaffolds, with *K*=200) against tag SNP sets from various genotyping arrays with sizes ranging from 30,710 to 49,191 tag SNPs in all populations. In this setting, fewer SNPs are used for LmTag compared to other arrays, as shown in Table 2. The compared arrays include widely used arrays include Genome-Wide Human SNP Array 6.0, Axiom Genome-Wide ASI, Axiom Genome-Wide EUR, Infinium Global Screening Array v3.0; and recently developed arrays such as Axiom Precision Medicine Diversity Array, Axiom Precision Medicine Research Array; and also customized-population-specific arrays including Axiom UK Biobank Array, Axiom Japonica Array NEO. Manifests of arrays are downloaded from respective manufacturers’ websites. Details of tested arrays and their corresponding number of tag SNP in chromosome 10 are reported in Table 2. Tag SNPs in chromosome 10 were then extracted and harmonized to the UCSC hg38 reference genome coordinate with CrossMap v0.2.6 if lifted over is required to obtain final tag SNP sets (Zhao et al., 2014). Imputation performances are estimated through leave-one-out cross-validation as described previously.

The comparison yields results as shown in Table 7, and Figure 6. In general, LmTag’s tag SNP sets outperform all compared array tag SNP sets. At 32,000 tag SNP scaffold, LmTag achieves 87.19%, 88.50%, 86.50%, and 89.80% overall imputation accuracies in EUR, EAS, SAS, and VNP, respectively, while the corresponding performances at 28,000 tag SNPs scaffold are 86.03%, 87.60%, 85.30%, and 88.91%. We also observe that population-specific optimization and size of the tag SNP sets in the arrays are two main factors affecting imputation performances. For instance, the recently developed Axiom Japonica Array NEO (Sakurai-Yageta et al., 2020) and the Axiom UK Biobank Array (Bycroft et al., 2018) performed best in the EAS and EUR populations with 84.70%, and 87.24% overall imputation accuracies, respectively. Besides, small size global optimization arrays such as the Infinium Global Screening Array v3.0 (30,710 tag SNPs in chromosome 10) shows the poorest performances across populations with 78.35%, 83.15%, 77.77%, and 82.81% overall imputation accuracies in EUR, EAS, SAS, and VNP, respectively. On the other hand, the Genome-Wide Human SNP Array 6.0 (49191 tag SNPs in chromosome 10) obtain much higher performances of 81.40%, 84.64%, 82.40%, and 85.69% for the same populations, respectively.

**Table 7:** Imputation performances of tag SNP sets selected by LmTag in examined populations (with *K* = 200 at 28000, and 32000 tag SNPs) against tag SNP sets extracted from various genotyping arrays. The imputation accuarcies are reported by various MAF bins from 0.01-0.5 and the “Overall” column reports mean imputation accuracy of all SNPs.

| Population | Array | 0.01:0.025 | 0.025:0.05 | 0.05:0.075 | 0.075:0.125 | 0.125:0.2 | 0.2:0.3 | 0.3:0.4 | 0.45:0.5 | Overall |
| --- | --- | --- | --- | --- | --- | --- | --- | --- | --- | --- |
| EAS | Affymetrix_6.0 | 56.21 | 74.06 | 83.51 | 87.77 | 89.23 | 89.46 | 89.15 | 88.56 | 81.40 |
| EAS | Axiom_GW_ASI | 55.86 | 73.63 | 82.73 | 86.76 | 88.53 | 88.21 | 87.51 | 87.43 | 80.48 |
| EAS | Axiom_GW_EUR | 52.98 | 70.49 | 80.73 | 84.65 | 87.12 | 86.68 | 86.28 | 86.11 | 78.53 |
| EAS | Axiom_JAPONICA | 62.17 | 77.97 | 87.96 | 90.86 | 91.55 | 91.52 | 91.17 | 91.29 | 84.70 |
| EAS | Axiom_PMDA | 53.55 | 72.19 | 80.85 | 85.19 | 86.75 | 86.95 | 86.06 | 86.12 | 78.85 |
| EAS | Axiom_PMRA | 59.69 | 75.06 | 82.66 | 85.77 | 86.82 | 87.18 | 86.62 | 86.23 | 80.52 |
| EAS | Axiom_UKB | 53.37 | 69.53 | 80.52 | 85.03 | 87.01 | 87.36 | 87.00 | 86.36 | 78.74 |
| EAS | Infinium_GSA | 56.07 | 72.71 | 80.16 | 83.67 | 85.54 | 85.50 | 85.02 | 84.18 | 78.35 |
| EAS | LmTag_K200_28000 | 67.44 | 81.51 | 88.06 | 90.72 | 91.62 | 91.77 | 91.16 | 91.17 | 86.03 |
| EAS | LmTag_K200_32000 | 70.25 | 83.10 | 89.10 | 91.48 | 92.23 | 92.39 | 91.85 | 91.85 | 87.19 |
| EUR | Affymetrix_6.0 | 66.48 | 79.04 | 86.71 | 88.77 | 91.06 | 91.28 | 91.09 | 91.61 | 84.64 |
| EUR | Axiom_GW_ASI | 63.99 | 76.23 | 84.20 | 86.94 | 88.88 | 89.56 | 89.16 | 89.51 | 82.43 |
| EUR | Axiom_GW_EUR | 67.11 | 80.53 | 87.42 | 89.75 | 90.27 | 90.64 | 90.08 | 90.30 | 84.68 |
| EUR | Axiom_JAPONICA | 65.46 | 77.35 | 84.56 | 85.99 | 89.89 | 90.57 | 90.90 | 91.10 | 83.41 |
| EUR | Axiom_PMDA | 68.37 | 80.44 | 86.90 | 88.69 | 89.88 | 89.99 | 89.82 | 89.97 | 84.54 |
| EUR | Axiom_PMRA | 67.11 | 78.84 | 85.56 | 87.66 | 88.99 | 89.08 | 88.62 | 89.34 | 83.38 |
| EUR | Axiom_UKB | 73.59 | 84.44 | 89.26 | 91.14 | 91.60 | 91.67 | 91.27 | 91.67 | 87.24 |
| EUR | Infinium_GSA | 69.47 | 80.90 | 83.29 | 85.79 | 87.47 | 88.11 | 88.22 | 87.74 | 83.15 |
| EUR | LmTag_K200_28000 | 72.73 | 83.55 | 89.64 | 91.66 | 92.76 | 92.66 | 92.48 | 92.13 | 87.60 |
| EUR | LmTag_K200_32000 | 74.37 | 84.63 | 90.42 | 92.23 | 93.43 | 93.38 | 93.17 | 92.80 | 88.50 |
| SAS | Affymetrix_6.0 | 63.94 | 77.37 | 84.40 | 88.59 | 89.38 | 90.26 | 90.17 | 90.68 | 82.40 |
| SAS | Axiom_GW_ASI | 61.29 | 74.33 | 81.74 | 86.29 | 87.46 | 88.38 | 87.68 | 88.74 | 80.02 |
| SAS | Axiom_GW_EUR | 62.74 | 74.83 | 82.22 | 86.53 | 87.44 | 88.36 | 87.66 | 88.89 | 80.44 |
| SAS | Axiom_JAPONICA | 62.18 | 75.12 | 82.51 | 87.64 | 88.81 | 90.07 | 90.16 | 90.82 | 81.42 |
| SAS | Axiom_PMDA | 63.18 | 76.23 | 83.79 | 86.68 | 87.38 | 87.80 | 87.66 | 87.80 | 80.69 |
| SAS | Axiom_PMRA | 61.89 | 74.34 | 81.99 | 85.57 | 86.35 | 87.14 | 86.75 | 87.24 | 79.51 |
| SAS | Axiom_UKB | 64.66 | 76.04 | 83.06 | 87.28 | 88.09 | 88.89 | 88.63 | 89.18 | 81.45 |
| SAS | Infinium_GSA | 59.74 | 72.21 | 80.34 | 83.72 | 84.61 | 85.45 | 85.55 | 86.06 | 77.77 |
| SAS | LmTag_K200_28000 | 70.44 | 81.71 | 88.35 | 90.71 | 91.12 | 91.04 | 90.76 | 91.07 | 85.30 |
| SAS | LmTag_K200_32000 | 72.39 | 83.21 | 89.41 | 91.54 | 92.03 | 91.91 | 91.74 | 92.01 | 86.50 |
| VNP | Affymetrix_6.0 | 70.26 | 79.70 | 86.00 | 89.20 | 90.61 | 90.90 | 90.66 | 91.00 | 85.69 |
| VNP | Axiom_GW_ASI | 69.33 | 78.69 | 84.88 | 88.18 | 89.94 | 89.78 | 89.19 | 89.79 | 84.65 |
| VNP | Axiom_GW_EUR | 67.25 | 76.51 | 83.28 | 86.35 | 88.52 | 88.63 | 87.91 | 88.74 | 83.07 |
| VNP | Axiom_JAPONICA | 73.85 | 81.59 | 88.93 | 91.48 | 92.52 | 92.60 | 92.21 | 92.90 | 87.89 |
| VNP | Axiom_PMDA | 66.59 | 77.45 | 83.32 | 86.93 | 88.36 | 88.69 | 87.98 | 88.55 | 83.11 |
| VNP | Axiom_PMRA | 70.35 | 79.30 | 84.79 | 87.29 | 88.53 | 88.83 | 88.33 | 88.57 | 84.18 |
| VNP | Axiom_UKB | 67.58 | 75.55 | 83.71 | 86.79 | 88.74 | 89.14 | 88.85 | 89.28 | 83.38 |
| VNP | Infinium_GSA | 68.12 | 78.15 | 83.44 | 85.98 | 87.45 | 87.62 | 87.28 | 87.64 | 82.81 |
| VNP | LmTag_K200_28000 | 77.61 | 84.91 | 89.18 | 91.35 | 92.44 | 92.83 | 92.39 | 92.66 | 88.91 |
| VNP | LmTag_K200_32000 | 79.68 | 86.13 | 90.08 | 91.93 | 92.99 | 93.31 | 93.01 | 93.21 | 89.80 |

**Figure 6:**
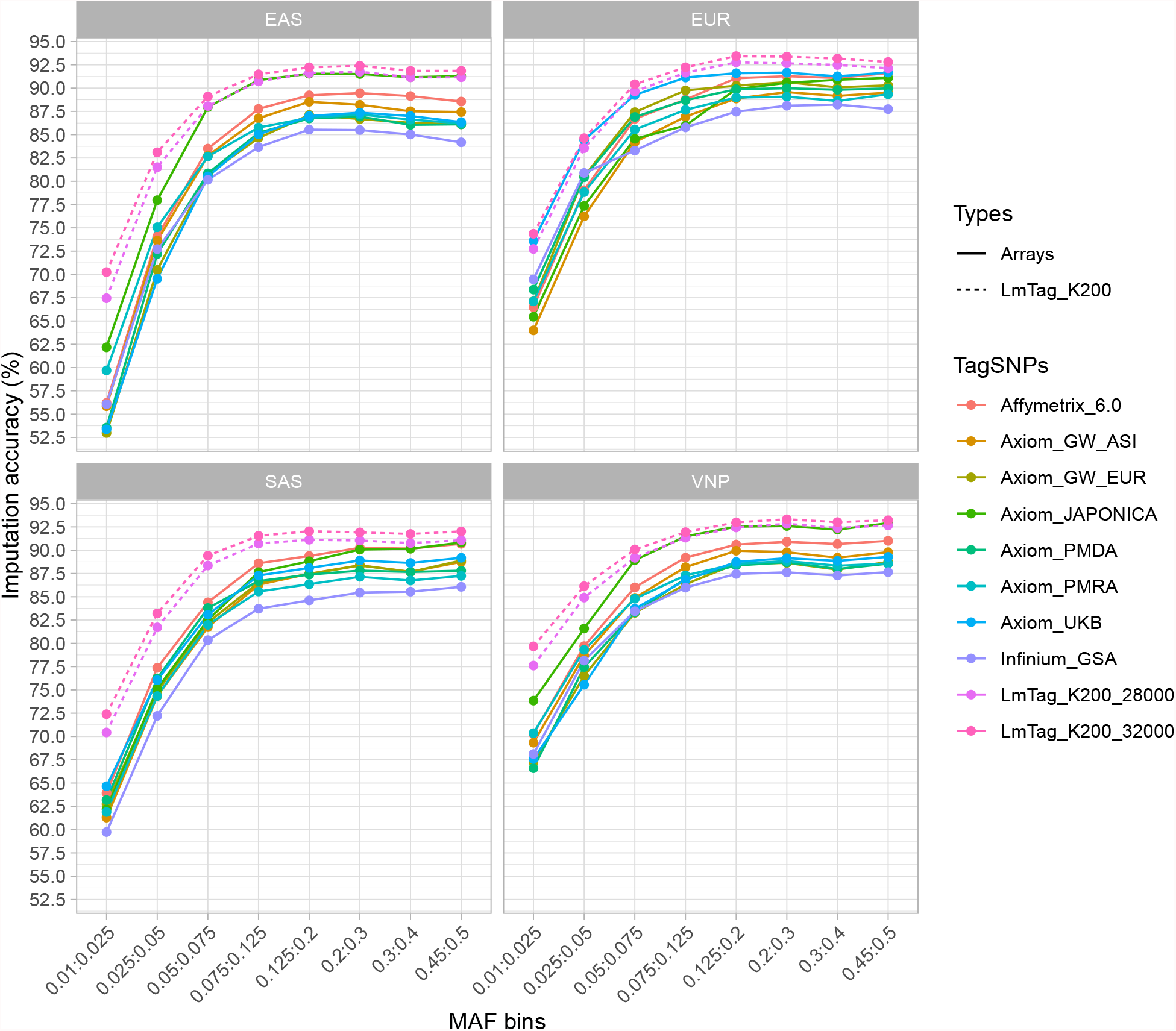
Imputation performances of various genotyping SNP arrays (Chromosome 10) in comparison with 28000 and 32000 tag SNP selected by LmTag (with *K* = 200).

Overall, LmTag can offer higher performance genotyping arrays with less number of tag SNPs compared to existing arrays. The imputation improvements vary from 9% compared to the Infinium Global Screening Array v3.0 in the EAS population to 1.5% compared to Axiom UK Biobank Array in the EUR population. Notably, for the VNP population, LmTag’s tag SNP sets specific for VNP appears to improve the imputation coverage the most compared to all other arrays, as shown in Table 7.

## Discussion and conclusions

Early genome-wide SNP arrays were usually designed by selecting tag SNPs from reference panels of predominantly European population (Rosenberg et al., 2010). As a result, these arrays often produce poorer performance in non-European populations (Altshuler et al., 2008; Rosenberg et al., 2010). Using customized, small-size SNP arrays at population-specific levels has recently emerged as an extremely cost-effective genotyping solution for underrepresented populations (Tam et al., 2019). For small arrays, imputation capability is essential to increase the genotyping coverage across the genome to capture as many DNA variants as possible. In addition to imputation performance, researchers also focus on the functional aspect of tag SNPs that are used in SNP arrays, which can help with fine mapping and increase the chance to detect true causal variants associated with traits. A recent comparative study of genotyping SNP arrays (Verlouw et al., 2021) discussed the importance of selecting markers based on biological-evidence and CADD functional scores (Rentzsch et al., 2019). In this study, we introduce a novel method, LmTag, that is optimized for both imputation and inclusion of functional variants. We compare the performance of LmTag to current widely used methods including EQ_uniform, EQ_MAF, TagIt, and FastTagger; and tag SNP sets from various SNP arrays. These methods and array designs are evaluated across four different populations. The results show that LmTag not only achieves higher imputation performance than other approaches but also significantly enrichs the tag SNP set with functional variants. Furthermore, results from our comparative analysis against existing SNP arrays suggest that LmTag has a high potential for designing new genotyping arrays, especially for underrepresented populations.

The improvement of tagging efficiency is mainly contributed by the LmTag statistical model. Instead of utilizing solely pairwise LD information as in conventional methods such as TagIt, LmTag assesses the relationship between imputation accuracy, mirror allele frequency, pairwise LD, and genomic distance and then uses this relationship to compute imputation scores for ranking SNP candidates tagging procedure. The model explains from 26.31% up to 44.15% imputation accuracy, depending on the genetic structure of populations. In all cases, the significant association of the model parameters with imputation accuracies are found, although the effect sizes varied across populations as shown in Table 8. While pairwise LD, MAF of both tag SNPs and tagged SNPs positively correlate with imputation accuracy, genomic distance showed the reverse trend.

**Table 8:** Estimated model parameters for relationship of imputation accuracy, minor allele frequency, pairwise LD, and genomic distance.

| Population | Intercept | LD $r^2$ | Tag SNP AF | Tagged SNP AF | Distance (MB) | Adjusted R-squared |
| --- | --- | --- | --- | --- | --- | --- |
| VNP | 0.64040*** | 0.30530*** | 0.00825*** | 0.01604*** | -0.13170*** | 42.76% |
| SAS | 0.61950*** | 0.32940*** | 0.01198*** | 0.01713*** | -0.21910*** | 40.79% |
| EUR | 0.66140*** | 0.28640*** | 0.02002*** | 0.01720*** | -0.20860*** | 38.26% |
| EAS | 0.55490*** | 0.38280*** | 0.01014*** | 0.02000*** | -0.16820*** | 44.15% |

\*\*\*: $p - value > 2e - 16$ .

Another advantage of LmTag is the implementation of beam search that considers a secondary factor in tag SNP selection. Besides genome-wide imputation capability, the inclusion of likely functional variants can enhance the value of genotyping SNP arrays by producing key information on potential causal SNPs underlying phenotypes. For example, the UK Biobank Axiom Array (Bycroft et al., 2018), Japonica NEO Arrays (Sakurai-Yageta et al., 2020), and the Axiom KoreanChip (Moon et al., 2019) applied various selection criteria to include likely functional markers in their array designs. However, these functional SNPs were selected independently from tag SNP selection procedure, i.e. no prioritization of tag SNPs regarding their biological functions was implemented. We introduce here an approach of searching for tag SNPs that are also highly functional. We believe that our proposed method will facilitate the next generation genotyping arrays that have high imputation performance as well as high biological functional potential that would facilitate post GWAS analysis such as statistical fine-mapping (Schaid et al., 2018) and the elucidation of biological mechanisms underlying the relationship between genotypes and phenotypes. Notably, in this study, we demonstrate how LmTag work in human datasets and CADD scores are used as a metric to approximate functional terms. Still, in practice, users could apply the method in other species with any criteria as long as they can provide a ranking scale for each SNP. For example, in other non-model species where calling confidence of the markers is a crucial factor, the method can be adapted for marker quality scores instead of functional scores, as long as a ranking system is provided.

## Methods

### Imputation accuracy modeling

Our aim is to combine systematical information from both pairwise LD *r*^2^, MAF, and genomic distance to improve imputation accuracy of tag SNP selection strategy. To this end, we model imputation accuracy as a linear model:

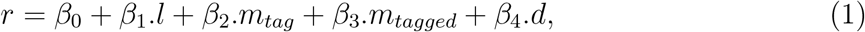

where:

1. *r* is imputation *r*^2^ (described later).
2. *l* is LD *r*^2^ between tag SNP and tagged SNP, (*l* ∈ (0 : 1]).
3. *m*_*tag*_ is MAF of tag SNP, (*m*_*tag*_ ∈ (0 : 0.5]).
4. *m*_*tagged*_ is MAF of tagged SNP (*m*_*tagged*_ ∈ (0 : 0.5])..
5. *d* is genomic distance between tag SNP and tagged SNP, (*d* ∈ *N*).

In this model, untyped SNPs are assumed to be tagged by the highest LD SNP in the tag SNP set. The relations among pairwise LD *r*^2^, MAF, and genomic distance are established by simulation. In details, a theoretical naive SNP array is created following by imputation accuracy scores computation for corresponding tagged SNPs. The corresponding information including LD, genomic distance are then extracted before using to estimate parameters for the linear model.

To make the simulation model as realistic as possible, we run a standard greedy tag SNP selection algorithm, TagIt (https://github.com/statgen/TagIt) (Weale et al., 2003), with default parameters (LD *r*^2^ threshold is 0.8, and MAF threshold is 0.01) to estimate scaffold sizes *k* for each chromosome. We denote the input containing *n* SNPs as *A* = {*SNP*_1_, *SNP*_2_, *SNP*_3_, …, *SNP*_*n*_}. We then sorted them by their genomic positions and uniformly sub-sampled *k* SNPs as a tag SNP set *T* = {*SNP*_1_, *SNP*_2_, …, *SNP*_*k*_}. The remaining *n* - *k* SNPs are labeled as a tagged SNP set *G* = {*SNP*_*k*+1_, *SNP*_*k*+2_, …, *SNP*_*n*_}. Imputation accuracy scores for tagged SNPs ∈ *G* are computed with a leave-one-out internal validation approach (Nelson et al., 2013; Wojcik et al., 2018). Specifically, imputation is performed individually for each sample with the exclusion of itself from the reference panel with Minimac4 v1.0.2 (Das et al., 2016). Tag SNPs ∈ *T* are denoted as ‘genotyped’ and the sites ∈ *G* are set as missing. The imputation accuracy *i*_*i*_ for each tagged *SNP*_*i*_ ∈ *G* is represented by the concordance rate, e.g., squared Pearson’s correlation coefficient which we termed imputation *r*^2^ to make a distinction from LD *r*^2^, between imputed genotype dosages in (0–2) and masked ground truth genotypes in (0, 1, 2).

Pairwise LDs are calculated using Plink v1.9 within a maximum genomic distance of 1 megabase (MB), and minimum LD *r*^2^ cutoff of 0.2 (Chang et al., 2015). Allele frequencies are computed and extracted with bcftools v1.10.2 (https://github.com/samtools/bcftools). To simplify the linear model, we assume that each tagged SNP’s genotype is inferred based on the sole tag SNP that has the highest LD *r*^2^. Thus, we find the best tag *SNP*_*i*_ ∈ *T* for each *SNP*_*j*_ ∈ *G* that has the most LD with the targeted tag *SNP*_*j*_ to extract relevant information including LD pairwise *l*_*ij*_, MAF *m*_*i*_, *m*_*j*_, and genomic distance *d*_*ij*_. Together with imputation scores *r*_*i*_ estimated from the previous step, these data are then used to estimate parameters for the linear model (1).

### SNP prioritization with high functional scores

In general, markers are functionally ranked based on biological evidence and genome-wide predicted functional scores. In the current implementation, GWAS catalog and the ClinVar databases are used to select biological SNP markers (MacArthur et al., 2017; Landrum et al., 2018). SNPs in these databases are functionally ranked to as in the highest score category. For non-biological evidence SNPs, we use Combined Annotation-Dependent Depletion (CADD) scores to prioritize functional SNPs (Kircher et al., 2014). The CADD scoring system is a widely used metric that effectively prioritizes causal variants in genetic analyses, especially in highly penetrant contributors to severe Mendelian disorders. CADD integrates more than 60 genomic features based on DNA sequence, for examples gene model annotations, evolutionary constraint, epigenetic measurements, and functional predictors into a single score by a machine learning model. In addition to the comprehensive use of genomic features, two other key advantages of the CADD model include the genome-wide estimation and the interpretability for each estimate. CADD scores are computed for all approximately 9 billion possible single nucleotide variants (SNV) across the human genome. For interpretability, the scores are transformed into ‘PHRED-scaled’ to provide a relative ranking system between SNVs at genome-wide coverage. Regardless of the details of the annotation set and model parameters, CADD scores can be interpreted simply as follows: a scaled score of 10 or greater equivalent to a raw score in the top 10% of all possible reference genome SNVs, and a score of 20 or greater indicates a raw score in the top 1%, and so on (Rentzsch et al., 2019).

### Functional tag SNP selection

Similar to most LD based tag SNP selection methods (Carlson et al., 2004; Weale et al., 2003; Liu et al., 2010; Hoffmann et al., 2011a,b), we employ a greedy approach for computational efficiency. However, there are two main differences in our algorithm. Firstly, we use estimated pairwise imputation *r*^2^ scores for ranking SNP candidates instead of using pairwise LD *r*^2^ like conventional methods. Specifically, for each pair of SNPs, imputation score *r*^2^ for each SNP is estimated independently by using coefficients derived from the established linear model and the corresponding LD *r*^2^, it’s MAF, mate’s MAF, and genomic distance between the two SNPs. Given two SNPs, *SNP*_*i*_, and *SNP*_*j*_ with LD *r*^2^ (*SNP*_*i*_, *SNP*_*j*_) = *l*_*ij*_, MAF *SNP*_*i*_ = *m*_*i*_, MAF *SNP*_*j*_ = *m*_*j*_, and genomic distance (*SNP*_*i*_, *SNP*_*j*_) = *d*_*ij*_. Their estimated imputation scores 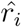, and 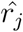 are calculated as:

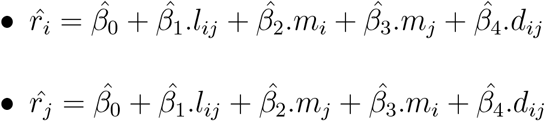

where 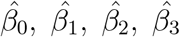, and 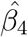 are estimated from the linear model (1). Secondly, LmTag employs beam search (Lowerre, 1976) instead of the best-first search strategy like other algorithms. The main advantages of beam search is allowing us to prioritize highly functional SNPs. In details, we introduce a tuning parameter *K* in the algorithm to select tag SNPs with high functional scores. LmTag algorithm starts with an empty tag SNP set *T*, a tagged SNP set *G*, and *n* input SNP candidates *A* = {*SNP*_1_, *SNP*_2_, *SNP*_3_, …, *SNP*_*n*_}. For each iteration, the algorithm includes two main steps as follows:

1. Imputation scoring. Each *SNP*_*i*_ ∈ *A* is scored as *s*_*i*_ which is sum of estimated imputation *r*^2^ 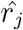 of all its neighboring *SNP*_*j*_ ∈ *A* given that pairwise LD *r*^2^ *l*_*ij*_ is equal to or greater than a specific cut-off *c*:

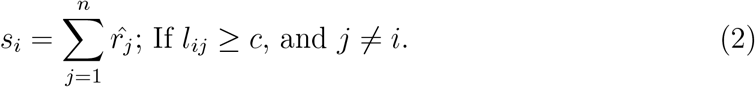
2. Tag SNP selection with beam search. Our approach considers the functional term of each marker in tag SNP selection by adapting the beam search algorithm (Lowerre, 1976). In brief, beam search is a heuristic searching algorithm used to solve combinatorial optimization problems. This approach employs a truncated branch-and-bound searching procedure where only the most promising *K* nodes (instead of all nodes) at each level of the search tree are evaluated and retained for further branching; *K* is the so-called beam width (Valente and Alves, 2005). We consider top *K* SNPs with highest imputation scores *s* as a candidate list of tag SNPs. Then, the search branching is extended to functional scores, i.e. the SNP with the highest functional score in this list is subsequently chosen as a tag *SNP*_*t*_. This SNP is subsequently moved from the candidate set *A* into tag SNP set *T*, and *SNP*_*t*_’s neighboring SNPs (satisfying pairwise LD *r*^2^ cut-off) are moved into tagged SNP set *G*. Overall, both the selected tag SNP and its associated tagged SNPs are removed from the candidate set *A*.

These steps are iterated until either *A* is empty or no pair (*SNP*_*i*_, *SNP*_*j*_) ∈ *A* satisfying the condition *l*_*ij*_ ≥ *c* could be found. Finally, the tag SNP set *A*, and their associated tagged SNP set *G* are exported.

### Datasets

The genomic data of the 1KVG pilot phase were obtained from 504 unrelated Vietnamese (Kinh ethnic group), including 208 males and 296 females. Their genomes were sequenced at coverage 30x with 150bp paired-end reads using an Illumina NovaSeq 6000 system. Variant calling was performed using the DRAGEN pipeline (Miller et al., 2015) with the GRCh38 patch release 13 reference genome (Van der Auwera et al., 2013). Quality check and filtering were performed with bcftools v1.10.2, and phasing was performed with SHAPEIT v4.1.3 to obtain the phased genotypes in Variant Call Format (VCF) (Delaneau et al., 2019).

Phased genotype data in VCF format of 1KGP NYGC high coverage are obtained from The International Genome Sample Resource (IGSR) data portal. We include only unrelated samples belonging to EAS, EUR, and SAS in the analysis. These samples are assigned to their super population according to IGSR’s annotation. All genomic data are reprocessed with bcftools v1.10.2 to keep only biallelic SNP with minor allele frequency (MAF) *>* 1%.

CADD v1.6 database (Rentzsch et al., 2019), release version 2021-07-08 of the GWAS catalog (MacArthur et al., 2017), and the ClinVar database (Landrum et al., 2018) are downloaded and filtered to obtain functional scores for each population. Finally, processed genomic data of four populations and their associated functional annotation are used in our analysis, including VNP, EAS, EUR, and SAS, which comprise 504, 504, 503, and 489 individuals respectively. Details of used datasets can be found in 1. Due to limited computational resources, our analysis was performed on chromosome 10, but the results obtained should be generalizable to all chromosomes.

### Performance evaluation

We compare LmTag against commonly used methods in SNP array design including TagIt (Weale et al., 2003), FastTagger (Liu et al., 2010), EQ_uniform (Shashkova et al., 2020), and EQ_MAF (Herry et al., 2018) using various metrics including imputation accuracy and functional enrichment. We also compare imputation accuracies of tag SNPs selected by LmTag against those of tag SNP sets from various commercial genotyping arrays. By this way, we want to explore potential applications of LmTag in designing genotyping arrays.

In terms of current methods for genotyping array design, the first two methods optimize imputation accuracy by maximizing linkage disequilibrium while the later methods select makers based on the equidistant principle. The distance-based methods are widely used in animal SNP array designs that involve dividing chromosomes into certain intervals with equal genomic length (Hayes et al., 2012; Shashkova et al., 2020; Joshi et al., 2018) and further optimized toward MAF (Dassonneville et al., 2012; Qiao et al., 2017; Herry et al., 2018). For each interval, the SNP with the highest MAF is selected as representative of all SNPs in the interval (Herry et al., 2018). TagIt is a typical greedy algorithm selecting tag SNPs based on pairwise LD information widely used in human SNP array designs (Weale et al., 2003; Sakurai-Yageta et al., 2020; Wojcik et al., 2018). Meanwhile, FastTagger is a fast implementation of the multi-marker LD approach, which reduces the number of tag SNPs selected while still maintaining high genomic coverage. In brief, the multi-marker LD approach methods find association rules of one SNP with multiple SNPs, termed multimarker *r*^2^ statistics, and use this information to find tag SNPs (Hao, 2007; Hao et al., 2007; Liu et al., 2010). Details on comparing these methods can be found in previous reports (Nguyen et al., 2021).

Evaluation metrics are based on imputation accuracy and functional prioritizing. Imputation accuracy is measured as squared Pearson’s correlation of imputed dosages estimated through a leave-one-out internal validation and the ‘true genotypes’. In details, selected tag SNPs are denoted as ‘genotyped’, and other sites are set as missing. For each SNP, squared Pearson’s correlation is calculated from imputation ‘estimated dosages’ (0-2) to the ‘true genotypes’ (0,1,2) in the original VCF file (Hoffmann et al., 2011a; Nelson et al., 2013; Wojcik et al., 2018). An overall imputation value is defined as mean imputation *r*^2^ of all markers in the population. Functional prioritizing is evaluated based on CADD scores and their corresponding percentiles among all SNPs, and the relative proportion of GWAS and ClinVar markers which is defined by the number of GWAS and ClinVar markers in the tag SNP sets over their corresponding number in the examined populations. These parameters are defined as follow:

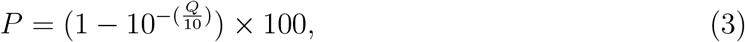

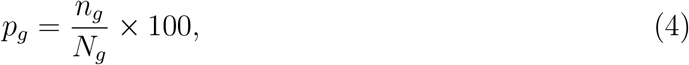

and

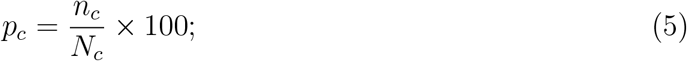

where *P* is percentile ranking of CADD score; *Q* is its original scores in ‘PHRED-scaled’; *N*_*g*_ and *N*_*c*_ are total GWAS and ClinVar markers in each populations; *n*_*g*_, *n*_*c*_ are number of GWAS and ClinVar markers in selected tag SNP sets; and *p*_*g*_, *p*_*c*_ are their corresponding proportions.

For comparison between methods, the LD cutoff is set at 0.8 in LD-based methods, including LmTag, TagIt, and FastTagger. FastTagger requires further LD settings for min_r2_2, and min_r2_3 that were set as 0.9, and 0.95 respectively, as recommended by the authors. LmTag is further ran with several *K* values varying from 1 to 200 to examine the relationship between imputation accuracy and functional SNP inclusion. Functional scores of selected tag SNPs by the other tag SNP selection methods are also computed for comparison. To enable a fair and comprehensive evaluation, tag SNPs are selected corresponding to multiple cutoffs ranging from 8000 to 32000 in all populations.

## Availability of data and materials

The 1KGP-NYGC datasets are freely available at IGSR data portal (https://www.internationalgenome.org). The 1KVG pilot phase datasets are available under agreement at MASH data portal (https://genome.vinbigdata.org/). LmTag is available for research only purpose at: https://github.com/datngu/LmTag. Data and source codes to generate figures of this study are available at: https://github.com/datngu/LmTag_data_analysis.

## Acknowledgements

We especially thank Nguyen T. Nguyen for his kindly help in downloading the 1KGP-NYGC datasets; Tran T.H. Tran, Mai H. Tran, and the 1000 Vietnamese Genomes Project team for their support in working with the VNP dataset; Khai Q. Tran for his useful suggestions in the C++ implementation; and Nghia T. Vu for his insightful comments during the early version of this manuscript. We also thank our colleagues in Center for Biomedical Informatics, Vingroup Big Data Institute, Vietnam and Institute for Molecular Bioscience, University of Queensland, Australia for their insightful discussions during the study.

## Authors’ contributions

DTN: conceptualized and implemented the algorithm, designed experiments, analyzed and interpreted results, drafted the manuscript. QHN and NSV: contributed to the discussion and manuscript revision. NTD and NSV: coordinated the project and supervised the study. All authors read and approved the final manuscript.

